# Schlafen 11 Restricts Flavivirus Replication

**DOI:** 10.1101/434563

**Authors:** Federico Valdez, Julienne Salvador, Pedro Palermo, Jonathon E. Mohl, Kathryn A. Hanley, Douglas Watts, Manuel Llano

**Author notes:** **Corresponding author:** Manuel Llano M.D., Ph.D., Department of Biological Sciences. University of Texas at El Paso. 500 West University, Ave. El Paso, TX 79968 USA.

## Abstract

Schlafen 11 (Slfn11) is a ubiquitously expressed interferon stimulating gene (ISG) that controls synthesis of proteins by regulating tRNA abundance. Likely through this mechanism, Slfn11 has previously been shown to impair human immunodeficiency virus 1 (HIV-1) infection and the expression of codon-biased open reading frames. Because replication of positive-sense single-stranded RNA [(+)ssRNA viruses] requires the immediate translation of the incoming viral genome whereas negative sense, single stranded [(−)ssRNA] viruses carry at infection an RNA replicase that makes multiple translation competent copies of the incoming viral genome, we reasoned that (+)ssRNA viruses will be more sensitive to the effect of Slfn11 on protein synthesis than (−)ssRNA viruses. To evaluate this hypothesis, we tested the effects of Slfn11 on the replication of a panel of ssRNA viruses in the human glioblastoma cell line A172, which naturally expresses Slfn11. Depletion of Slfn11 in this cell line significantly increased the replication of (+)ssRNA viruses from the *Flavivirus family*, including West Nile (WNV), dengue (DENV), and Zika virus (ZIKV) but had no significant effect on the replication of the (−)ssRNA viruses vesicular stomatitis (VSV, *Rhabdoviridae family*) and Rift Valley fever (RVFV, *Phenuiviridae family*). Despite that WNV titers in Slfn11-deficient cells were almost 100-fold higher than in cells expressing this protein; they produced approximately two-fold less viral particles, as determined by PCR-based quantification of virion-associated WNV RNA in the cell culture supernatant. These data indicated that Slfn11 impairs WNV fitness but does not affect other steps of the viral life cycle including entry, viral RNA replication and translation, and budding. Similarly to the proposed anti-HIV-1 mechanism of Slfn11, this protein prevented WNV-induced down-regulation of a subset of tRNAs implicated in the translation of 19% of the viral polyprotein. Importantly, we provided evidence suggesting that the broad anti-viral activity of Slfn11 requires other cellular proteins, since overexpression of Slfn11 in cells that naturally lack the expression of this protein, did not impair WNV or HIV-1 infection. In summary, this study demonstrates that Slfn11 restricts flaviviruses replication by impairing viral fitness.

**AUTHOR SUMMARY:** The host targets mechanisms that viruses have evolved to optimize replication. We provide evidence that the cellular protein Schlafen 11 (Slf11) impairs replication of flaviviruses, including West Nile (WNV), dengue (DENV), and Zika virus (ZIKV). However, replication of single-stranded, negative RNA viruses was not affected. Specifically, Slf11 decreases the fitness of WNV potentially by preventing virus-induced modifications of the host tRNA repertoire that could lead to enhanced viral protein folding. Furthermore, we demonstrated that Slf11 is not the limiting factor of this novel broad anti-viral pathway.

## INTRODUCTION

Successful viral replication depends on the ability of the virus to appropriate the host translational machinery. The innate immune response exploits this dependency to control viral replication. Many interferon (IFN)-stimulated genes (ISGs) that regulate protein translation are well known to restrict virus replication, including Protein Kinase R, the Interferon-induced proteins with tetratricopeptide repeats family of proteins, zinc-finger antiviral protein and the 2′,5′-Oligoadenylate/RNaseL pathway. Another family of ISGs is the Schlafen (Slfn) proteins, which were first identified as being important regulators of T cell differentiation and growth[1, 2]. Currently, 10 mouse (Slfn1, 1L, 2, 3, 5, 8, 9, 10, and 14) and 6 human (Slfn5, 11, 12, 12L, 13, and 14) Slfn genes have been identified[1, 2]. Slfn11, the focus of the current study, is ubiquitously expressed in human tissues[3] but is absent in mice[1, 2]. This protein controls synthesis of proteins encoded by codon-biased open reading frames[3–5].

Several members of the Schlafen family have been shown to impair virus replication. Mouse Slfn14 impairs replication of influenza A and varicella zoster virus. The mechanism for this effect is unknown but Slfn14 affects nuclear trafficking of influenza nucleoproteins and enhances IFN-β signaling[6]. Human Slfn11 suppresses HIV-1 and equine infectious anemia virus infection[4, 5]. Slfn11 binds to tRNAs and counteracts the up-regulation of the tRNA repertoire induced by HIV-1 infection that promotes translation of the codon-biased viral genome[4]. The anti-viral mechanism of Slfn11 seems to involve a tRNA nucleolytic activity recently described in Slfn13[7]. Eight out of nine residues implicated in the tRNA nucleolytic activity of Slfn13[7] are conserved in Slfn11, and these two proteins share an overall homology of 83%. This enzymatic activity is required for Slfn13 to restrict HIV-1 infection. Slfn13 cleaves tRNAs close to the 3’ end at the acceptor stem and also diminishes the levels of HIV-1 mRNA. The anti-HIV-1 activity of Slf13 is specific since this protein did not affect replication of herpes simplex or Zika viruses (ZIKV)[7].

Considering the effect of Slfn11 on protein synthesis, we hypothesized that this protein will preferentially restrict the replication of positive-sense single-stranded RNA [(+)ssRNA] viruses over (−)ssRNA viruses. Replication of (+)ssRNA viruses requires the immediate translation of the incoming viral genome. In contrast, (−)ssRNA viruses do not have this dependency as they introduce upon infection a RNA-dependent RNA polymerase that uses the incoming viral genome as template to produce multiple copies of translation competent viral RNA. Therefore, (+)ssRNA viruses are expected to be more sensitive to impaired protein translation than (−)ssRNA viruses, as recently evidenced[8].

Based on this, we predicted that knockdown of Slfn11 would enhance replication of (+)ssRNA viruses but have no effect on replication of (−)ssRNA viruses. Consequently, we predicted that overexpression of Slfn11 would restrict replication of (+)ssRNA but not (−)ssRNA viruses. Finally, we expected to see that WNV infection will modulate the tRNA repertoire in Slfn11-deficient cells but not in cells expressing this protein. We tested these predictions and found that flaviviruses, including West Nile (WNV), dengue (DENV), and ZIKV, replicated significantly more efficiently in Slfn11-deficient than in control cells expressing this protein. However, this phenotype was not observed when the replication of (−)ssRNA viruses vesicular stomatitis (VSV, *Rhabdoviridae* family) and Rift Valley fever (RVFV, *Phenuiviridae* family) was analyzed, highlighting the specificity of this anti-viral activity. Furthermore, we demonstrated by PCR-based quantification of virion-associated WNV RNA in the cell culture supernatant that although WNV titers were almost 100-fold higher in Slfn11-deficient cells than in cells expressing this protein; the deficient cells produced two-fold less viral particles, indicating that Slfn11 impairs WNV fitness. These data also excluded an affect of Slfn11 on entry, viral RNA replication and translation, and budding. Analysis of the tRNA repertoire indicated that Slfn11 prevented WNV-induced down-regulated the expression of a subset of tRNAs implicated in the translation of the viral polyprotein Finally, we found that cells lacking endogenous Slfn11 failed to support the anti-viral activity of exogenously introduced Slfn11. In summary, our data demonstrated that Slfn11 decrease viral fitness restricting replication of Flaviviruses.

## RESULTS

### WNV infection induces expression of Slfn11

To determine whether WNV infection modulates levels of Slf11 expression, cells of the human glioblastoma cell line A172 were infected with WNV at MOI: 0.1 and viral replication and expression of Slfn11 was determined at different times post-infection (p.i.) by plaque assay and immunoblot, respectively (Fig. 1a-b). Viral replication was detectable as early as 8 hrs p.i. and peaked by 32 hrs after infection (Fig. 1a). Corresponding with the peak of viral replication, we detected a sustained increase in the basal levels of Slfn11 after 40 hrs p.i. (Fig. 1b). Densitometry analysis of immunoblots from two independent infection experiments indicated that WNV infection caused 3.5-fold increase in the α-tubulin-normalized Slfn11 protein levels after 40 hrs p.i. Therefore, these data indicated that Slfn11 is up-regulated by WNV infection.

**Figure 1.**
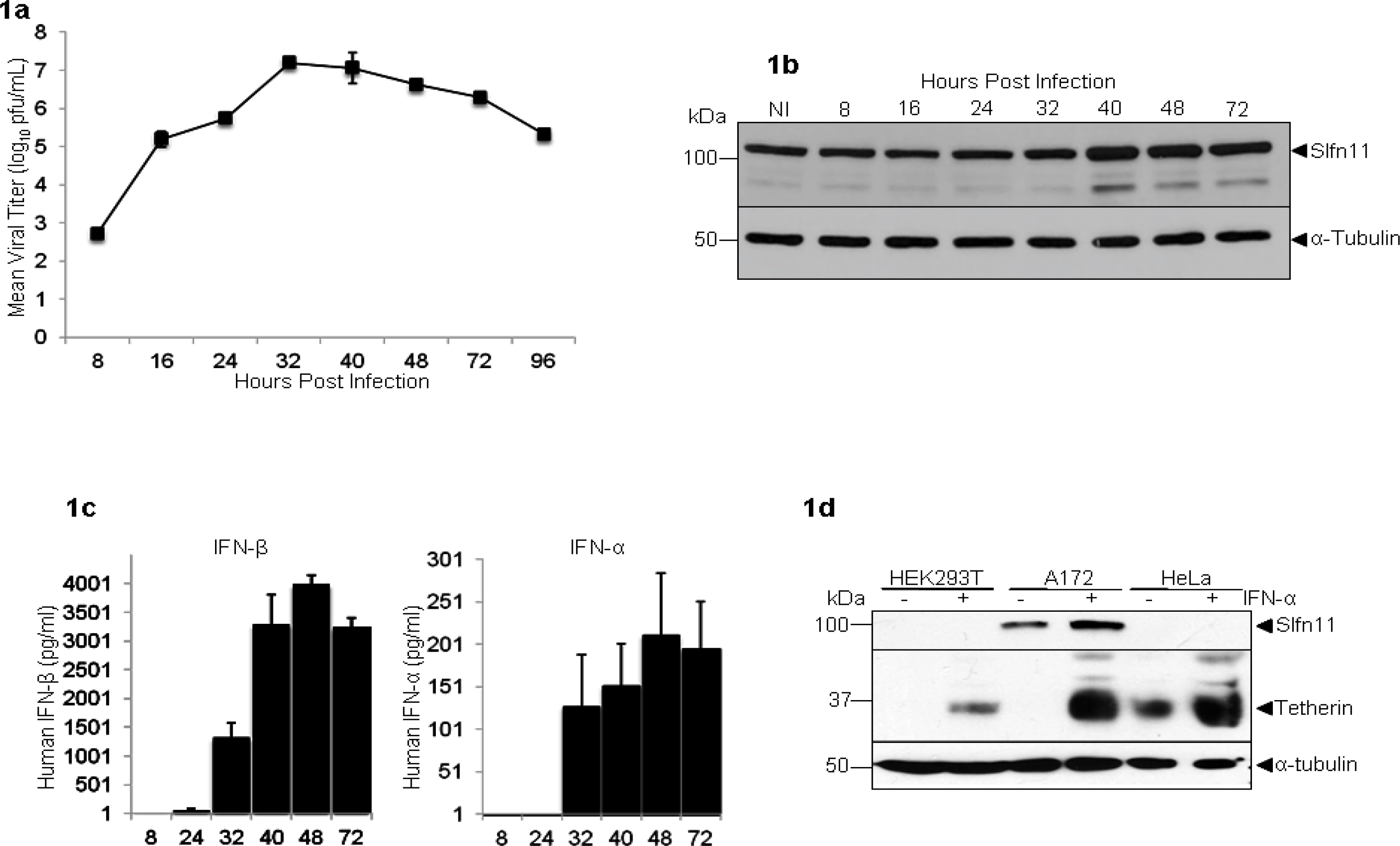
Kinetics of WNV replication, type I interferon production, and Slfn11 expression in A172 cells. (**a**) WNV replication in A172 cells. Cells were infected with WNV at MOI 0.1 and viral replication was measured by titration of the cell supernatant in a plaque assay at different times post-infection. Titers were determined in triplicate experiments. Data are representative of 3 independent infection experiments. (**b**) Expression of Slfn11 in WNV-infected A172 cells. Cells were lysed at different times post-infection and Slfn11 and α-tubulin (loading control) were detected with specific antibodies by immunoblot. Data are representative of 2 independent infection experiments. (**c**) Kinetics of IFN-α and IFN-β (all subtypes) production in WNV-infected A172 cells. Culture supernatant was collected at different times post-infection and type I IFN quantified by ELISA. Data are representative of 3 independent infection experiments. (**d**). Effect of IFN-α1 on Slfn11 expression. HeLa, A172, and HEK293T cells were treated with 5000U/ml units of IFN-α1 for 24 hrs and the expression of the type I IFN-stimulated genes Slfn11 and Tetherin were evaluated by immunoblot.

The up-regulation of Slfn11 in WNV-infected cells could be secondary to the production of type I IFN in response to the viral infection. Therefore, we evaluated the temporal sequence in production of these proteins. A172 cells were infected with WNV, as described above and levels of IFN-α and −β were determined in the cell supernatant by ELISA. IFN-α and −β were undetectable, <1.95 pg/ml and <2.3pg/ml, respectively, at 8 hrs p.i. even though viral replication was evident by this time (Fig. 1c). However, type I IFNs production was evident by 32 hrs, reaching a peak at 48 hrs after infection. Therefore, these data indicated a temporal correspondence between type I IFN secretion and the up-regulation of Slfn11 suggesting that virus-induced type I IFN up-regulated Slfn11 expression.

To further evaluate the role of type I IFN in the regulation of Slfn11 expression a panel of cell lines susceptible to WNV infection was treated with IFN α-1 for 24 hrs and then the levels of Slfn11 was determined by immunoblot. Similar to WNV infection, IFN α-1 triggered a two-fold increase of Slfn11 in A172 cells (Fig. 1d). However, this treatment failed to induce expression of Slfn11 in HEK293T or HeLa cells, which also lack basal expression of this protein. To verify that IFN α-1 stimulated these cells, we also measured the expression of the ISG tetherin. As previously reported, tetherin was constitutively expressed in HeLa cells but not in HEK293T cells and the expression of this protein increased in both cell types in response to IFN α-1 stimulation (Fig. 1d). Tetherin was absent in untreated A172 cells but was also significantly induced after IFN α-1 treatment. Therefore, the lack of response to IFN α-1 was not the reason for the absence of Slfn11 in HeLa and HEK293T cells.

### Slfn11 impairs replication of flaviviruses but not of (−)ssRNA viruses

To test the relative effect of Slfn 11 expression on (+)ssRNA and (−)ssRNA viruses, A172 cells were stably transduced with lentiviral vectors expressing shRNAs containing Slfn11-specific[4] or scrambled sequences to generate Slfn11-deficient A172 cells (A172-KD) and control cells (A172-SCR), respectively. Subsequently, A172-KD cells were engineered to over-express Slfn11 (A172-BC). Levels of Slfn11 were verified in these cells by immunoblotting (Fig. 2a). Then, A172-derived cell lines were infected with WNV at a MOI of 0.1 and cell culture supernatants were collected every 8 hrs to measure viral titer by plaque assay.

**Figure 2.**
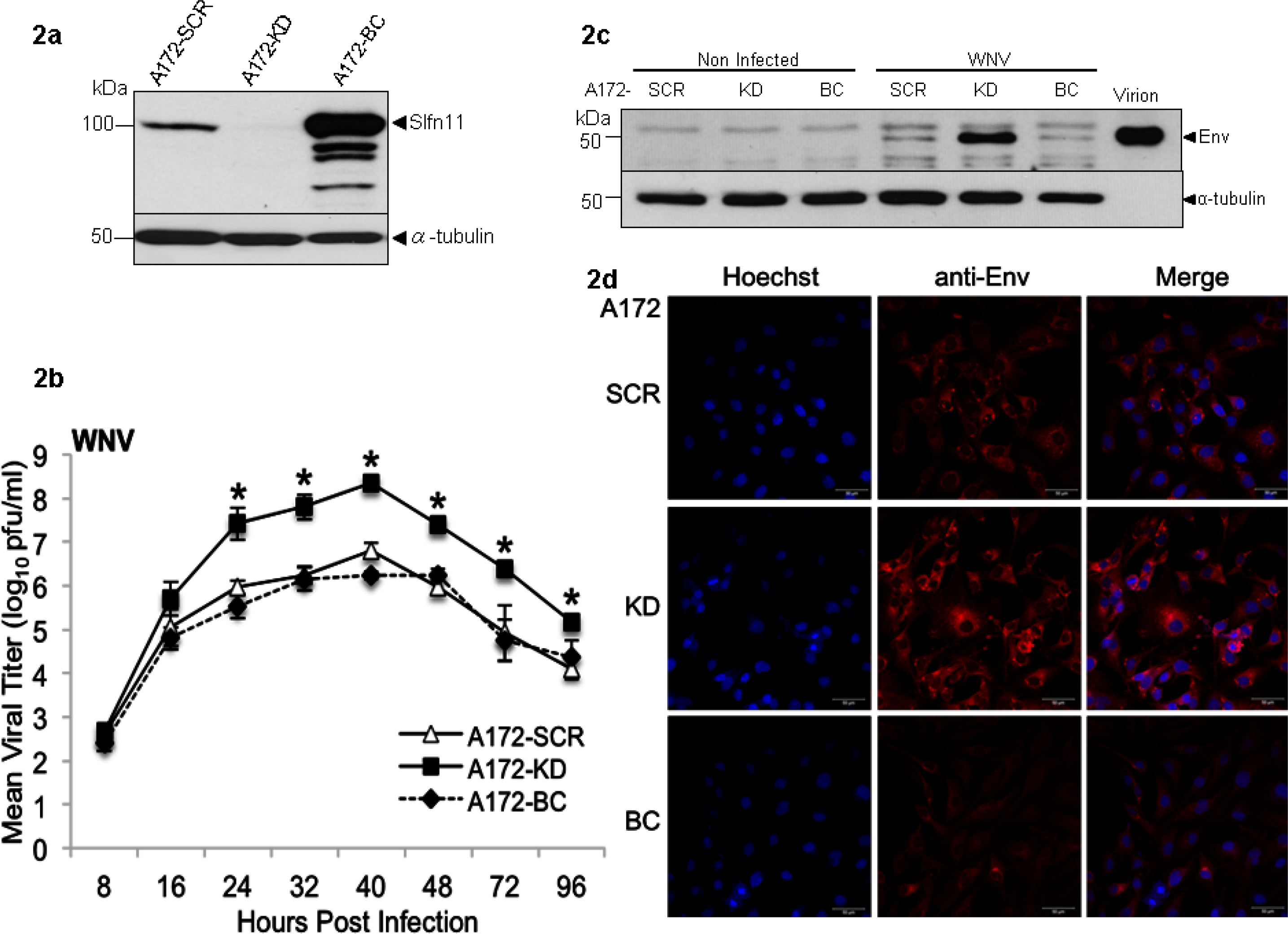
Effect of Slfn11 on WNV replication. (**a**) Immunoblot analysis of the expression of Slfn11 in A172 cells stably expressing shRNAs directed against Slfn11 (A172-KD) or a scrambled (A172-SCR) RNA sequences and A172-KD cells engineered to re-express Slfn11 (A172-BC). α-tubulin was detected as a loading control. (**b**) WNV replication in A172-derived cells. A172-SCR (open triangles), A172-KD (filled diamonds) and A172-BC (filled squares) cells were infected with WNV (MOI 0.1) and viral replication was determined by quantification of the viral titer in the cell supernatant at different hours post-infection by plaque assay. Statistically significant differences were calculated with repeated measures ANOVA and Tukey-Kramer post-hoc tests and they are indicated with asterisks. Mean and standard deviation values represent the variability of the viral titer found in triplicate plaque assays of samples from 8 independent infection experiments performed in different days with different viral preparations. (**c**) Expression of WNV Env in cells infected at MOI 0.1 evaluated by immunoblot 40 hrs after infection. α-tubulin was detected as a loading control. These results are representative of 3 independent infections. (**d**) Expression of WNV Env (red) as detected by indirect immunofluorescence analysis of cells infected at a MOI of 1 48 hrs post-infection. Nuclei were labeled with Hoechst (blue).

For all the replication curves reported in this study, we analyzed the data with repeated measures ANOVA, and we always detected a significant effect of time p.i. on virus titer; as this effect was expected and is not of interest we do not report it here. Instead we focus on whether there was a significant interaction between treatment (i.e. cell type) and time, or, in the absence of such interaction, an independent effect of treatment, and we use Tukey-Kramer post-hoc tests to elucidate the nature of these effects.

As shown in Figure 2b, there was a significant interaction of the effects of cell type (KD, SCR or BC) and time p.i. on WNV titer (DF =14, F = 11.1, P< 0.0001). Overall, WNV replicated significantly more efficiently in A172 cells lacking Slfn11 than in either of the two cell lines expressing this protein (Fig 2b). Twenty-four hrs post- infection viral titers were 2 logs higher in Slfn11-deficient cells than in control cells and these differences persisted until 96 hrs post-infection, after which the experiments were terminated due to significant cytopathic effects.

We anticipated Slfn11 to impair WNV replication by targeting viral protein production. Therefore, we infected A172-derived cells with WNV at a MOI of 0.1 and measured cell-associated WNV Env at the peak of infection (40 hrs p.i.) by immunobloting analysis. As expected, Slfn11-deficient cells expressed higher levels of WNV Env than control cells (Fig. 2c). Densitometry analysis of the immunoblots corresponding to three independent infection experiments indicated that α-tubulin-normalized Env levels were similar between A172-SCR and −BC cells and A172-KD cells expressed 12.3-fold more Env protein than A172-SCR cells. These results were expected based on the WNV titer reached in these cells and in the postulated mechanism of action of Slfn11.

In addition, we verified WNV Env expression by indirect immunofluorescence. A172-derived cell lines were infected with WNV at MOI of 1 and the production of WNV Env was evaluated 24 and 48 hrs after infection by indirect immunofluorescence with an anti-flavivirus antibody that reacts with WNV Env. In correspondence with data in figure 2c, Slfn11-deficient cells expressed higher levels of Env than A172-SCR and −BC cells at 48 hrs p.i. (Fig. 2d). As expected, analysis at 24 hrs post-infection showed similar results (data not shown). Subcellular distribution of Env was similar in all the cell lines, indicating the absence of gross defects in Env intracellular trafficking. Therefore, data in figures 2b-d demonstrated that Slfn11 markedly impaired the replication of WNV by impairing viral protein expression.

We next tested whether Slfn11 affected replication of two additional flaviviruses, DENV and ZIKV. Contrary to WNV, DENV does not replicate efficiently in A172 cells; however, we were interested in determining the contribution of Slfn11 to this phenotype. Then, A172-derived cells were infected with DENV at MOI 0.1 and viral replication was followed at different times post-infection by titration of the cell supernatant by plaque assay. Data in figure 3a shown statistically significant interactions (DF=8, F=24.4, P<0.0001). Interestingly, although A172-SCR cells did not support replication of DENV (Fig. 3a), Slfn11-deficient A172 cells efficiently allowed DENV replication, producing 2-log higher viral titters than A172-SCR cells at the peak of replication. DENV reached titers of 10^6^ pfu/ml in A172-KD cells with a replication kinetic similar to that of WNV. In these experiments, DENV replication peaked from 24 to 32 hrs post-infection and then decayed by 48 hrs due to cytopathic effects. As expected, re-expression of Slfn11 in A172-KD cells (A172-BC cells) fully removed the permissiveness of these cells, highlighting the specificity of the effect of Slfn11 on DENV replication. Similarly to A172-SCR cells, DENV did not multiply in A172-BC cells. Therefore, basal expression levels of Slfn11 significantly contribute to the restriction of DENV replication in A172 cells.

**Figure 3.**
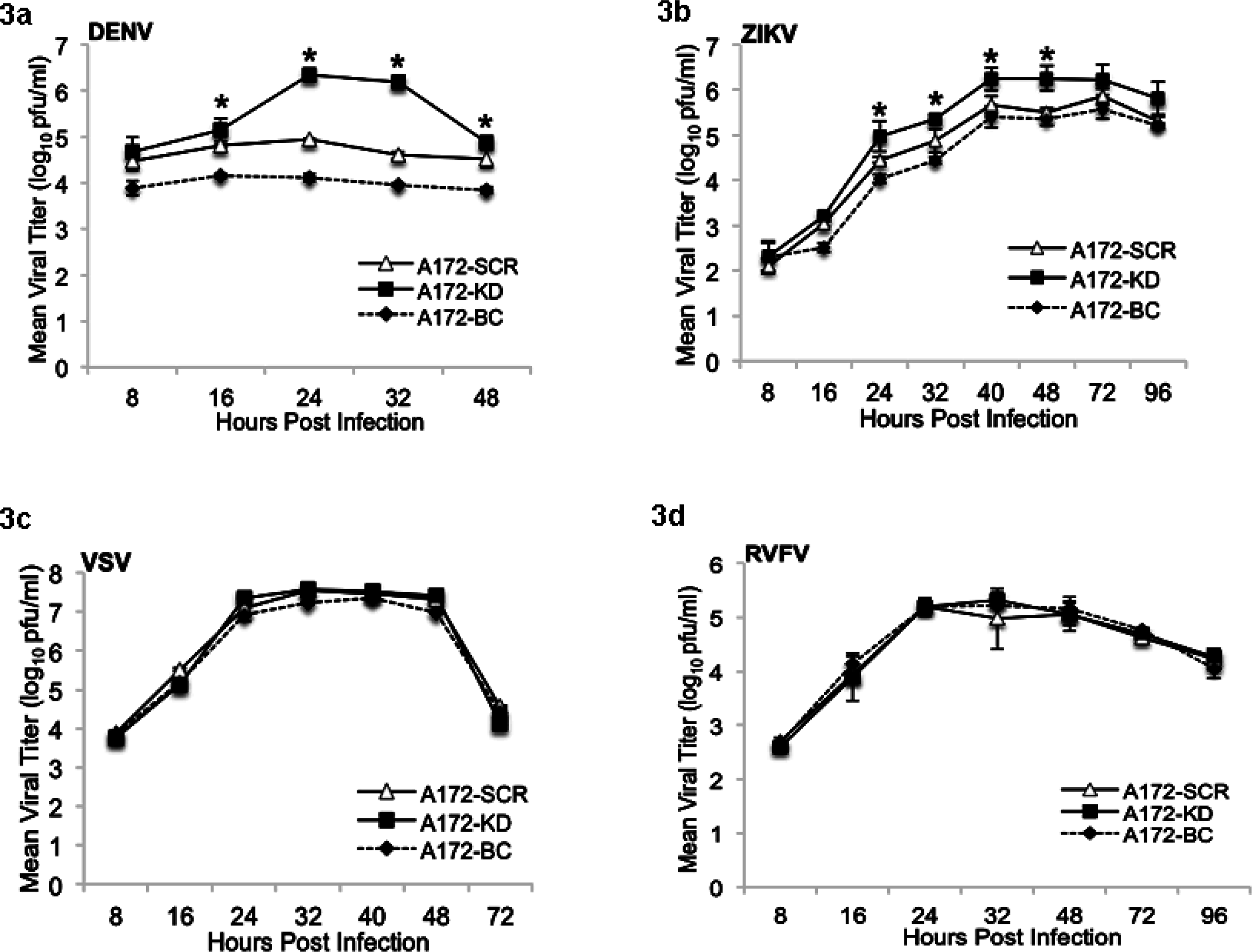
Effect of Slfn11 on viral replication. A172-SCR (open triangles), A172-KD (filled diamonds) and A172-BC (filled squares) cells were infected with (**a**) Dengue Virus (DENV, MOI 0.1), (**b**) Zika Virus (ZIKV, MOI 0.1), (**d**) Vesicular stomatitis virus (VSV, MOI 0.1), and (**e**) Rift Valley fever virus (RVFV, MOI 0.1). Viral replication was determined by quantification of the viral titer in the cell supernatant at different hours post-infection by plaque assay. Statistically significant differences are indicated with asterisks and were calculated as described above. Mean and standard deviation values of each graphic represent the variability of the viral titer found in triplicate plaque assays of samples from 3 independent infection experiments performed in different days with different viral preparations.

Slfn11 and Slfn13 impair HIV-1 infection through a similar mechanism[4, 7]. However, Slfn13 fails to restrict ZIKV replication[7] and the effect of Slfn11 on this flavivirus has not been explored yet. Thus, we determined whether or not Slfn11 impairs the replication of ZIKV. A172-derived cells were infected and ZIKV titer was quantified as described above. ZIKV replication peaked at 40 hrs post-infection and plateaued until 96 hrs post-infection or the end of the experiment. However, cytopathic effects were not very apparent even at these late time points of the infection. Importantly, replication of ZIKV was also significantly enhanced by deficiency of Slfn11 (Fig. 3b), although the intensity of the effect was less marked than for WNV and DENV (DF=14, F=3.44, P<0.0009). Nevertheless, a significant difference of approximately 7-fold higher titer was observed in A172-KD versus the control cells from 24 to 48 hrs post-infection. Therefore, Slfn11 restricts ZIKV replication in contrast to Slfn13[7].

For comparison with patterns of (+) and (−)ssRNA virus replication, we also tested the impact of Slfn11 knockdown and overexpression on the replication of the (−)ssRNA viruses VSV and MP12-RVFV. Replication of both viruses was very robust in A172 cells showing a kinetic similar to that of WNV with a peak of viral replication 24 hrs post-infection (Fig. 3c-d). However, the replication of these (−)ssRNA viruses was very similar in terms of kinetics and titers among the different A172-derived cell lines evaluated, despite their differences in Slfn11 expression. Therefore, there was not interaction between the effects of cell line and time p.i. on virus titer (VSV, DF=12, F=0.21, P=0.9963; RVFV, DF=12, F=0.74, P=0.6974).

The VSV used in these experiments is a recombinant virus expressing eGFP[9]. Therefore, we also evaluated the titer of this virus obtained in the supernatant of the different A172-derived cell lines by flow cytometry analysis. A172-SCR and −KD cells were infected with three different MOIs of VSV (0.1, 0.3, and 1) in triplicates and viral supernatant was collected 24 hrs later. Then, SupT1 cells were infected with the viral supernatants and evaluated 24 hrs after infection by FACS analysis. In concordance with the plaque assay experiments, we observed a similar titer for the virus obtained from A172-SCR (8.55 ×10_4_ +/− 0.84) and −KD (8.58 ×10_4_ +/− 0.91) cells. In summary, data in Figs 3c-d indicated that Slfn11 did not influence the replication of the (−)ssRNA VSV and RVFV, highlighting the specificity of the anti-viral activity of this protein for (+)ssRNA viruses and lentiviruses[4].

From the results presented about it is also noteworthy that despite A172-BC cells express markedly higher levels of Slfn11 than A172-SCR cells (Fig. 2a) replication of flaviviruses was similar in these two cell lines (Figs. 2b and 3a-b), indicating that above certain levels the anti-viral effect of Slfn11 reaches saturation.

### Mutagenesis analysis of Slfn11 anti-viral activity

The N-terminal region of Slfn11 and Slfn13 harbors the anti-HIV-1 activity of these proteins[7]. Not surprisingly, this region contains in both proteins the residues that in Slfn13 are responsible for the tRNA nucleolytic activity[7] which is central in the anti-HIV-1 activity of these proteins^3,13^. Therefore, we followed a previously described strategy [4] to determine whether the anti-WNV activity of Slfn11 also resides in the N-terminal region.

The N-terminal (amino acids 1-441) and the C-terminal (amino acids 442-901) regions of Slfn11 were expressed in A172-KD cells to generate A172-N-term and A172-C-term cell lines (Fig. 4a) and their susceptibility to WNV was evaluated (Fig. 4b). Cells were infected with WNV at MOI 0.1 and viral replication was followed by plaque assay, as described above. WNV replication was impaired in cells expressing N-terminal Slfn11. Viral replication was similar in these cells and in A172-BC cells that express full-length Slfn11. However, WNV replication was significantly higher in cells expressing Slfn11 C-terminus (DF=14, F=11.69, P<0.0001, Fig. 4b). Therefore, these findings suggest a common anti-viral mechanism of action of Slfn11 against HIV-1[4] and WNV.

**Figure 4.**
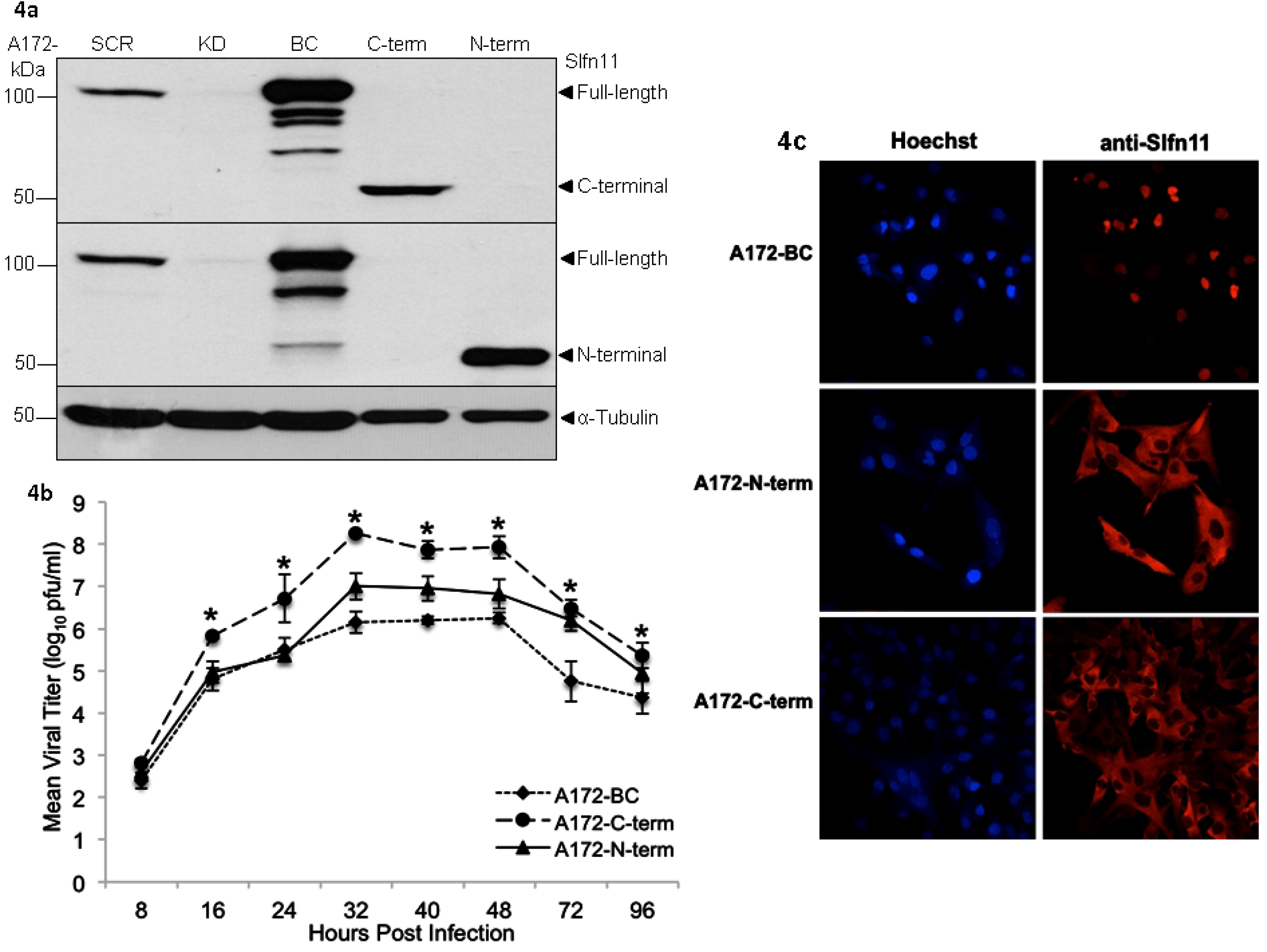
Mutagenesis analysis of anti-viral activity of Slfn11. (**a**) Immunoblotting analysis of the expression of Slfn11 in A172-derived cells. A172-KD cells were engineered to express the N- or C-terminus of Slfn11. A172-SCR, −KD, and BC were used as control. Different anti-Slfn11 antibodies were employed to identify the mutant proteins. α-tubulin was detected as a loading control. (**b**) A172-BC (filled diamonds), A172-N-term (filled triangles), and A172-C-term (filled circles) were infected with WNV (MOI 0.1) and viral replication was determined by quantification of the viral titer in the cell supernatant at different hours post-infection. Statistically significant differences were calculated as described above and are indicated with asterisks. Mean and standard deviation values represent the variability of the viral titer found in triplicate plaque assays of samples from 3 independent infection experiments. (**c**) Cellular distribution of Slfn11 full-length and deletion mutants. A172-BC, A172-N-term, and A172-C-term cells were fixed/permeabilized and stained with the anti-Slfn11 antibodies used in panel (**a**). Cell nuclei were identified with Hoechst staining (Blue staining).

In order to define whether the subcellular distribution impacts the anti-viral activity of Slfn11, we determined the localization of Slfn11 full-length and truncation mutants. A172-BC, −N-term and −C-term cells were stained with anti-Slfn11 antibodies directed against the N-terminus or the C-terminus of the protein and Slfn11 cellular distribution was determined by confocal microscopy analysis. Full-length Slfn11 accumulates in the nucleus, distributing homogenously in this organelle (Fig. 4c). However, both the C- and N-terminal Slfn11 proteins were uniformly distributed in the cytoplasm of the cell (Fig. 4c). The lack of association of anti-viral activity and subcellular distribution of Slfn11 suggests that the process targeted by this protein is accessible in both the nucleus and the cytosol. In addition, these data confirm that the inactivity of the C-terminal region of Slfn11 is not determined by miss localization or a gross defect in intracellular solubility of this protein, for example due to the formation of protein aggregates.

### Effect of WNV infection and Slfn11 expression on the tRNA repertoire of A172 cells

It has been previously reported that Slfn11 counteracts HIV-1-induced increase of tRNA abundance, affecting viral protein expression[4]. Therefore, we explored whether WNV infection also induces changes in the tRNA repertoire and whether these changes are opposed by Slfn11.

A172-SCR, −KD, and −BC cells were infected with WNV at a MOI of 1 and their tRNA repertoire was determined 8 hrs later by tRNA PCR array, as previously described[4, 10]. To ensure data robustness, the tRNA repertoire of A172-derived cells infected and non-infected was determined by using RNA obtained from three sets of independent experiments. We chose to evaluate the tRNA repertoire at 8 hrs post-infection because at this early time point of the life cycle the WNV was already replicating (Fig. 1a and 2b) and the infected cells exhibit similar viral loads (Fig. 2b), were not producing type I IFN (Fig. 1c) nor undergoing cell death (data not shown). Therefore, modifications in the tRNA pool at 8 hrs post-infection is likely be a direct consequence of the infection of these cells with similar amount of virus. Furthermore, at 8 hrs post-infection we expected WNV replication to be very sensitive to the efficiency of translation due to the low availability of translation competent viral RNA molecules caused by the low rate of synthesis of viral RNA characteristic of the early life cycle[11].

Comparison of the levels of 66 nuclear tRNAs in infected and non-infected A172-KD cells indicated that WNV infection decreased the expression of 10 tRNAs while 7 were up-regulated (Fig. 5a). The limited effect of WNV on the tRNA pool contrasted with to the global up-regulation of tRNAs triggered by HIV-1 in cells expressing low levels of Slfn11[4]. Out of these 17 tRNAs, 9 were down-regulated and 1 was up-regulated statistically significant by WNV infection in A172-KD cells (Fig. 5b), and none of these 10 tRNAs were modified in either in A172-SCR or −BC when infected cells and non-infected cells were compared. Then, these WNV-induced tRNA changes were considered Slfn11-specific (Fig. 5b).

**Figure 5.**
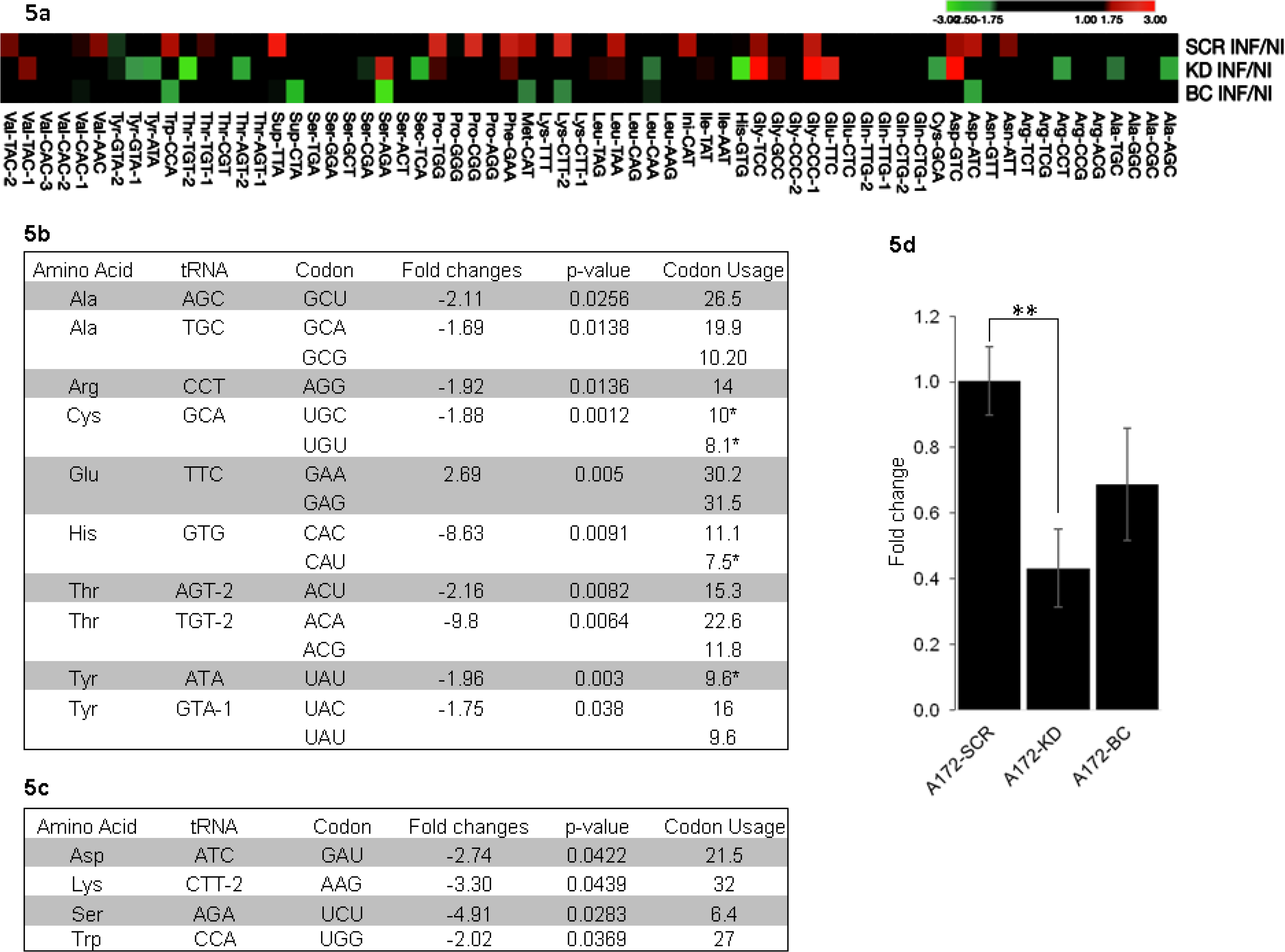
Effect of WNV infection on the tRNA pool of Slfn11-deficient and control cells. (**a**) Heat map of fold changes of the 66 tRNA quantified. Each cell represents the tRNA ratio of infected/non-infected cells for each of the three cell lines evaluated. tRNAs that increase (red cells), decrease (green cells) or do not change (black cells) are indicated. The cut-off was set at −1.75 to 1.75 fold change. tRNAs modulated in A172-KD cells by WNV infection. (**b**) WNV-induced, Slfn11-specific tRNA changes. Fold differences represents the A172-KD infected/non-infected tRNA ratio. (**c**) WNV-induced tRNA changes in A172-BC cells. Fold differences represents the A172-KD infected/non-infected tRNA ratio. In (**b**) and (**c**) the statistical significance of the changes in tRNA levels is represented by the calculated p values. The percentage of codons decoded by each tRNA is indicated. Rare codons are marked with an asterisk. **(d)**Virion-associated WNV RNA was quantified by reverse transcription-PCR in the in cell culture supernatant of A172-derived cells at 24 hrs post-infection (MOI 0.1). The viral titter of theses samples is represented in figure 2b. Results show the relative fold differences in virion-associated WNV RNA compared to the A172-SCR, control cell line. Mean and standard deviation values represent the variability of 2^-Ct values found in 3 independent infection experiments. Statistical analysis was performed by One-way ANOVA with Tukey HSD post-hoc test. ** Indicates p<0.01.

Notably, the tRNAs down-regulated by WNV in Slfn11-deficient cells (Fig. 5b) were implicated in decoding 19.2% of the viral codons, while those found up-regulated corresponded only to 7.7%. Analysis of the impact of tRNA changes on codons that WNV uses disproportionately more than human cells (rare codons, 12.3 % of the WNV codons) indicated that the cognate tRNA of 28.5 % of them were decreased by WNV infection in A172-KD cells, representing 3.5% of all the WNV codons. The codons affected by tRNA reductions were randomly distributed along the viral genome.

WNV infection did not consistently down- or up-regulated any of the 66 tRNAs analyzed in both A172-SCR and −BC cells (Fig. 5a). Instead, we noticed that tRNAs have a tendency to be up-regulated in A172-SCR and down-regulated in A172-BC following WNV infection. In A172-SCR cells 17 tRNAs increased; however, in A172-BC eight of them did not change and the other four decreased (Fig. 5a). This inconsistency suggest that these changes were independent of Slfn11. Alternatively; it could reflect a higher Slfn11 tRNA nucleolytic activity in A172-BC than in A172-SCR cells, as the engineered cells express more Slfn11 than A172-SCR cells (Fig. 2a). Nevertheless, WNV replicated similarly in A172-SCR and −BC cells, indicating that these changes in tRNA abundance did not impact WNV replication. Furthermore, analysis of the impact of tRNA down-regulation in A172-BC on the WNV polyprotein indicated that tRNAs in this group that exhibit statistically significant changes decode only common codons that represent 8.7% of the WNV genome (Fig. 5c).

WNV infection also modulated the expression of tRNAs that reprogram stop codons to encode specific amino acids (Fig. 5a). Genetic code reprogramming allows translation beyond stop codons producing fusion proteins[12]. WNV infection decreased the levels of the tRNA^Sec^(TCA) in A172-KD cells but not in cells expressing Slfn11, indicating that this change was Slfn11-specific. This tRNA incorporates selenocysteine in selenoproteins at the UGA stop codon. Furthermore, WNV infection diminished the expression of tRNA^Sup^(CTA) in A172-BC cells that redefines the stop codon UAG (Amber suppressor); however, this change was not observed in A172-SCR cells that also express Slfn11.

In conclusion, the changes in the tRNA repertoire inflicted by WNV infection suggested that the virus will encounter more restrictions to protein translation in Slfn11-deficient cells than in cells expressing this protein. This seems contradictory with the more robust WNV replication observed in these cells. However, multiple lines of evidence (discussed later) indicate that a reduced availability of cognate tRNAs pauses translation at elongation favoring either protein production and/or optimal protein folding[13–21]. The later mechanism enhances viral fitness [13, 15, 20].

In order to evaluate this hypothesis, we quantified by reverse transcription (RT)-PCR the viral load in the supernatant of cells expressing or not Slfn11. Viral supernatants harvested 24 hrs post-infection (analyzed in Fig. 2b) were subjected to RNase treatment to remove virion-free RNA and then RNA was isolated and used in RT-PCR with WNV-specific primers. In these experiments (Fig. 5d) we observed that A172-SCR and −BC cells produced 2.5− and 1.7-fold, respectively, more viral particles than A172-KD cells, despite that viral titters were near 100-fold higher in Slfn11-deficient cells than in those expressing this protein. These findings indicate that Slfn11 impairs viral fitness. Furthermore, these results exclude any negative effect of Slfn11 on other steps of the viral life cycle including entry, viral gene expression, and budding.

### Lack of anti-viral activity of Slfn11 in HEK293T, HeLa, and BHK1 cells

Slfn11 has been shown to mediate anti-viral activity in cells expressing endogenous Slfn11 such as CEM, HEK293 [3, 4], and A172 cells (Fig. 2b). However, the anti-WNV activity of this protein has not been evaluated in cells that naturally lack the expression of Slfn11 under basal conditions and/or following type I IFN stimulation (Fig. 1d). Among them, we characterized the WNV-permissive cell lines HEK293T, HeLa, and baby hamster kidney fibroblasts (BHK-21). These cell lines were engineered to express Slfn11 (HEK293T^Slfn11^, HeLa^Slfn11^, and BHK-21^Slfn11^ cells) (Fig. 6a) and then used in viral infection experiments.

**Figure 6.**
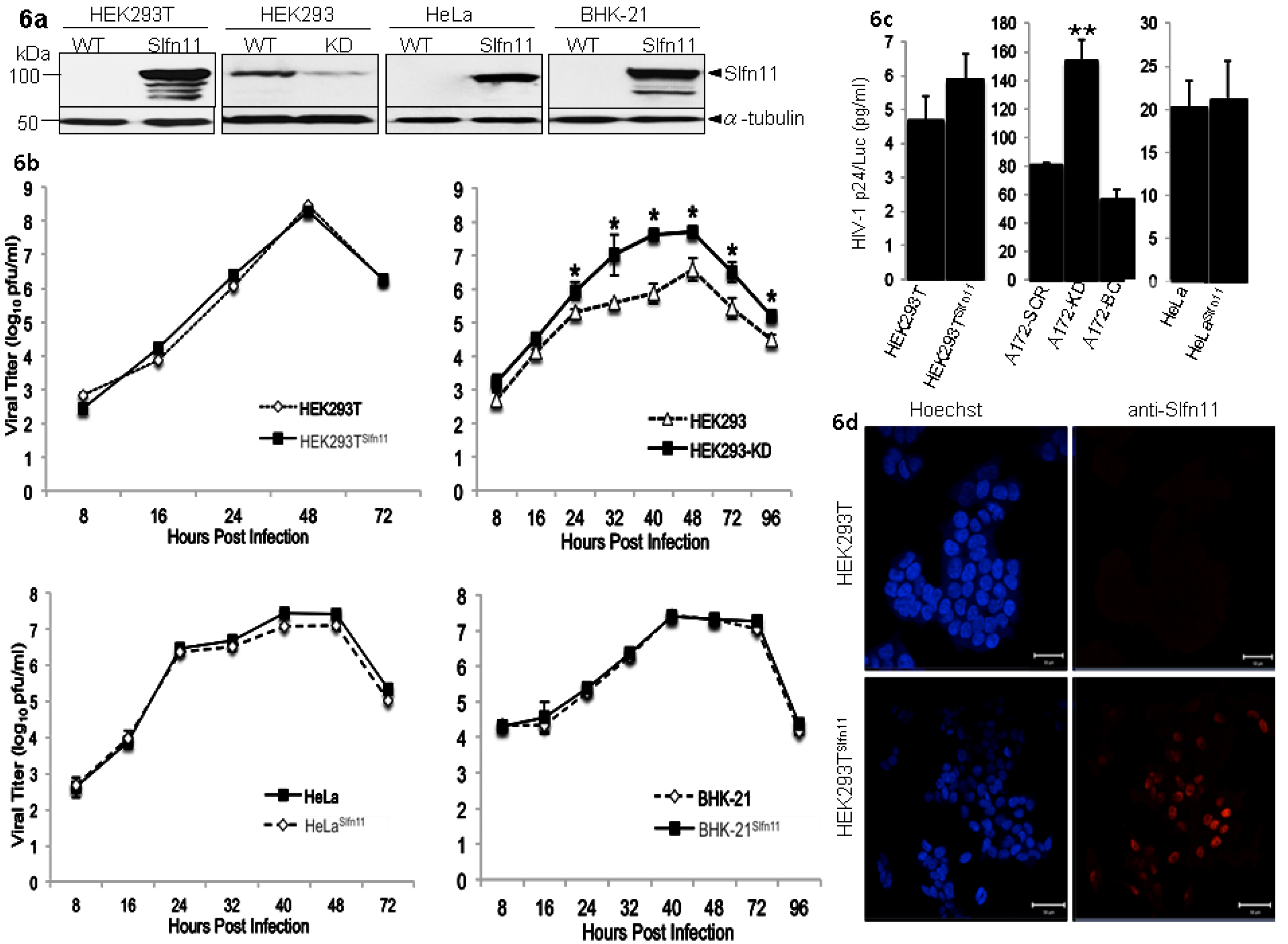
Evaluation of the anti-viral activity of Slfn11 in HEK293T, HeLa, and BHK-21 cells. (**a**) Slfn11 expression in HEK293T, HEK293, HeLa, and BHK-21 parental and derived cell lines as detected by immunoblotting analysis. (**b**) WNV replication in HEK293TSlfn11, HEK293-KD, HeLaSlfn11, and BHK-21Slfn11 and parental cell lines. Parental (open diamonds) and Slfn11-expressing derivative cell lines (filled triangles) and HEK293 (open triangle) and HEK293-KD (filled squares) cells were infected with WNV (MOI 0.1) and viral replication was determined by quantification of the viral titer in the cell supernatant at different hours post-infection by plaque assay. Statistically significant differences were calculated as described above and are indicated with asterisks. Mean and standard deviation values represent the variability of the viral titers found in triplicate plaque assays of samples from 3 independent infection experiments. (**c**) HIV-1 infection of cells expressing or not Slfn11. Cells were infected with a replication defective HIV-1 that expresses LTR-driven luciferase and p24. Mean and standard deviation values represent the variability of luciferase-normalized HIV-1 p24 levels found in 3 independent infection experiments. Statistical analysis was performed by t-test (HEK293 and HeLa) and ANOVA with Tukey HSD post-hoc test (A172). ** Indicates p<0.01 (**d**) Cellular distribution of Slfn11 in HEK293T^Slfn11^. Cells were fixed/permeabilized and stained with an anti-Slfn11 antibody by indirect immunofluorescence (red). Nuclei were labeled with Hoechst (blue).

HEK293T-derived cells were infected with WNV at MOI 0.1 and viral replication was determined by plaque assay as described above. WNV replicated robustly in these cells (Fig. 6b) with a kinetic similar to the replication in A172 cells (Fig. 1a). However, WNV replication was indistinguishable between HEK293T cells expressing or not Slfn11 (Fig. 6b).

To further corroborate the lack of activity of Slfn11 in cells that do not naturally produce it, we evaluated the susceptibility of HEK293T and HEK293T^Slfn11^ cells to a single-round infection HIV-1-derived reporter virus (Hluc). This virus expresses p24 and luciferase proteins from the viral promoter[22]. Slfn11 has been reported to affect expression of codon-biased but not unbiased open reading frames[3–5]; therefore, we expect that HIV-1 p24, but not luciferase activity, will be affected by expression of Slfn11 in HEK293T cells.

HEK293T and HEK293T^Slfn11^ cells were infected with Hluc and luciferase activity and p24 levels were measured in cell lysates and in the cell culture supernatant, respectively, 4 days after infection. As expected, we did not observe any significant effect of Slfn11 on luciferase expression. Cells expressing Slfn11 exhibited 1,740.4 +/− 121 arbitrary units (AU)/ml compared to parental cells that produced 1,188.4+/− 94.2 AU/ml. However, both luciferase-normalized (Fig. 6c) and non-normalized (data not shown) HIV-1 p24 levels were not affected by Slfn11 expression in HEK293T cells.

In contrast to HEK293T cells, the parental cell line HEK293 expresses endogenous Slfn11 and supports the anti-HIV-1 activity of this protein[4]. Therefore, we determined the effect of Slfn11 on WNV replication in HEK293. Control and Slfn11-deficient cells (Fig. 6a) were infected with WNV at a MOI of 0.1 and viral replication was followed by plaque assay as described above. WNV replication was very robust in HEK293 cells; however, Slfn11 knockdown significantly enhanced viral replication (DF=7, F=8.11, P<0.0001, Fig. 6b). These results demonstrated the anti-WNV activity of Slfn11 in HEK293 cells.

As an additional control, we evaluated the effect of Slfn11 on HIV-1 in A172-derived cells. In disparity with HEK293T cells, A172 cells express endogenous Slfn11 (Fig. 1d). A172-SCR, −KD, and −BC cells were infected with Hluc and four days later luciferase activity was measured in cell lysates and the expression of HIV-1 p24 in the cell supernatant. In these experiments we found similar levels of luciferase activity among the different cell lines evaluated indicating that Slfn11 levels did not affect the expression of a codon unbiased HIV-driven gene. A172-SCR cells produced 1,072 +/− 15 AU/ml and A172-KD cells 1.227.5+/− 6.1 AU/ml, whereas A172-BC cells exhibited 1,289.4 +/− 11.1 AU/ml of luciferase activity. In contrast to HEK293T cells, luciferase-normalized (Fig. 6c) and non-normalized (data not shown) HIV-1 p24 expression was 2-fold higher in A172-KD cells than in −SCR and −BC cells. Thus, these results were in agreement with previously reported observations in CEM and HEK293 cells[4], indicating that Slfn11 restrict HIV-1 in A172 cells. Furthermore, results in figure 6c indicated that Slfn11 similarly impaired HIV-1 infection in A172-SCR and −BC cells despite of their differences in Slfn11 expression levels (Fig. 2a). These findings are in agreement with the saturating effect that we described above for the anti-flavivirus activity of Slfn11 (Fig. 2), further supporting our hypothesis that Slfn11 is not the limiting factor of the anti-viral pathway.

We hypothesized that the absence of anti-viral activity of Slfn11 in HEK293T cells could be due to mislocalization of the exogenously expressed protein. Thus, we determined the subcellular distribution of Slfn11 in HEK293T^Slfn11^ cells by immunofluorescence analysis as described above. Similar to A172 cells (Fig. 3c), Slfn11 was localized entirely in the nucleus of HEK293T^Slfn11^ cells and protein aggregates were not observed (Fig. 6d). Therefore these findings exclude mislocalization or protein aggregation as a cause for the lack of activity of Slfn11 in HEK293T cells.

Next, we explored the effect of Slfn11 on WNV replication in HeLa and BHK-21 cells following the procedures described above. HeLa and BHK-21 parental and cells engineered to express high levels of Slfn11 (HeLa^Slfn11^ and BHK-21^Slfn11^ cells, Fig. 6a) were infected with WNV at MOI 0.1 and viral replication was followed by plaque assay. Similarly to HEK293T cells, replication of WNV in HeLa and BHK-21 was very robust and exhibited a kinetic comparable to the replication in A172 cells (Fig. 6b). However, in contrast to A172 cells, we did not find any differences in viral replication in HeLa or BHK-21 cells expressing or not Slfn11 (HeLa: DF=6, F=2.42, P=0.0562; BHK-21: DF=7, F=0.65, P=0.711; Fig. 6b).

Furthermore, we determined the effects of Slfn11 on single-round HIV-1 infection in HeLa cells as described earlier. Similarly to the previous experiments we did not observe differences in the luciferase levels of HeLa cells expressing or not Slf11. Parental cells expressed 18.73 +/− 2.63 AU/ml while cells engineered to express Slfn11 have 16.97 +/− 1.59 AU/ml. Importantly, HIV-1 p24 levels were also similar in the HeLa cell lines either prior or after luciferase normalization (Fig. 6c), indicating that Slfn11 did not impair HIV-1 p24 production in HeLa cells. Therefore, our data indicated that HEK293T, BHK-21, and HeLa cells do not support the anti-viral activity of exogenously expressed Slfn11. These results further support our hypothesis that Slfn11 is not the only component of this anti-viral pathway.

## DISCUSSION

Slfn11 and other members of this family, such as Slfn13 and Slfn14, have been shown to exhibit anti-viral functions[4, 6, 7]. In particular, Slfn11 impairs lentivirus infection, including HIV-1[3, 4] and Equine Infectious Anemia Virus[5]. Mechanistically, this protein blocks HIV-1-induced up-regulation of tRNA, impairing viral protein translation[4]. Slfn11 is absent in mice and the ortholog of human Slfn11 in this specie is unknown[1, 2], limiting the *in vivo* characterization of this protein.

The efficiency of protein translation is importantly influenced by tRNA availability; therefore, we postulated that Slfn11 will also selectively affect the replication of (+)ssRNA over (−)ssRNA viruses. In contrast to the latter group, (+)ssRNA viruses require the immediate translation of the incoming viral genome, making them more susceptible to impaired protein translation. Our data support these predictions. We found that Slfn11 severely affects replication of flaviviruses, including WNV, DENV, and ZIKV; however, the (−)ssRNA viruses VSV and RVFV were not affected. Therefore, the anti-viral activity of Slfn11 seems to be specific for (+)ssRNA viruses and lentiviruses[4, 5].

Our findings also revealed important differences between Slfn11 and Slfn13 in their specificity. Slfn13 failed to affect ZIKV replication[7] whereas we found Slfn11 effective against this virus. These functional differences highlight the lack of redundancy in the anti-viral specificity of different members of the Slfn family. Importantly, human placenta and testes lack expression of Slfn11, otherwise, a ubiquitously expressed protein[3]. We predict that the lack of expression of Slfn11 in these tissues could significantly influence mother to fetus and sex-related transmission of ZIKV[23–25].

Combined analysis of ours and previous findings[4] demonstrated that Slfn11 impairs HIV-1 and WNV through a similarly mechanism. In both cases the N-terminal region of Slfn11 is necessary and sufficient for anti-viral activity and this protein blocks virus-induced changes in the tRNA repertoire of the infected cells. However, HIV-1 globally increased tRNAs in the absence of Slfn11[4]; in contrast WNV infection only decreased a subset of tRNAs in Slfn11-deficient cells. These findings suggest that Slfn11 opposes virus-induced changes in the tRNA pool independently of their direction. In the case of HIV-1-infected cells, it is likely that Slfn11 decreases tRNA levels by degradation, similarly to Slfn13. Eight out of nine residues implicated in the tRNA nucleolytic activity of Slfn13[7] are conserved in Slfn11. However, we do not understand how Slfn11 counteract WNV-induced decrease in tRNA levels.

Ours results here and previously published data[4] suggest that lentiviruses and flaviviruses have evolved different strategies to enhance viral replication by modulating the tRNA pool. Lentivirus increases tRNA levels in the cell resulting in a higher availability of tRNAs required for the translation of their codon-biased genome. Therefore, these changes in the tRNA pool lead to enhanced viral protein translation[4, 26]. In contrast, flaviviruses, which lack codon-biased genomes, diminish the abundance of a subset of tRNAs predictably reducing the efficiency of viral protein production, and nonetheless, increasing virus replication.

Multiple lines of evidence indicate that pausing of translation elongation at codons recognized by low abundance tRNAs might promote efficiency of protein production and/or optimal protein folding [13, 15-21, 27]. A common pattern of 30-50 rare codons clustering immediately downstream the start codon is present in highly expressed prokaryotic and eukaryotic genes. It is though that this “ramp” sequence increases protein synthesis efficiency by reducing ribosome stalling during translational elongation [18]. However, it is unlikely that the WNV-induced tRNA decrease will enhance viral replication by this mechanism as the codons affected do not cluster in the viral genome.

Furthermore, it has been proposed that rare codons decrease the speed of translational elongation potentially facilitating protein folding [27]. In the RNA replicase and 3C protease of foot-and-mouth disease virus [15, 20] and in a *Neurospora crassa* circadian clock protein, a link between location of rare codons and protein secondary structures was described[21]. This link was also predicted by bioinformatics analysis of *E. coli*, *S. cerevisiae*, *Caenorhabditis elegans*, and *Drosophila melanogaster* genomes[21]. The location of these codons in the transition boundaries of protein secondary structures suggests that the modulation of the speed of the translation elongation is necessary for proper protein folding[27]. Furthermore, replacement of native rare codons by synonymous common codons in the capsid of poliovirus[28] and hepatitis A virus [13] caused a decrease in viral fitness, suggesting that optimal protein folding due to low abundance of tRNA results in increased viral fitness. Therefore, we postulate that WNV-induced reduction in tRNA abundance in Slfn11-deficient cells leads to optimal viral protein folding and enhanced viral fitness.

Our data further verified these predictions. Using quantitative reverse transcription-PCR of virion-associated RNA we found that, despite the titer of WNV produced in Slfn11-deficient cells was near to 100-fold higher than the titers of viruses produced in cells expressing Slfn11, the later produced close to twice more viral particles that the knockdown cells. These findings strongly suggest that Slfn11 impair WNV replicative fitness and that it does not have a negative effect on other steps of the viral life cycle.

Considering the lack of response of (−)ssRNA viruses VSV and RVFV to Slfn11, we propose that this type of viruses do not alter the abundance of the tRNA pool. However, we lack evidence to support this hypothesis because the effect of viral infection on the host tRNA composition is ill-defined. Nevertheless, it was demonstrated that influenza virus does not change the abundance of the tRNA repertoire[29], lending support to our hypothesis.

Intriguingly, we observed that exogenously expressed Slfn11 is not functional against HIV-1 or WNV infection in HEK293T, HeLa, and BHK1 cells that naturally lack endogenous production of this protein. These observations suggest that Slfn11 is not the only component of this broad anti-viral pathway. These findings also indicate that the anti-viral activity of Slfn11 is different from the effects of Slfn11 on the expression of transiently transfected plasmids [3], since the later activity was verified in HEK293T cells expressing exogenous Slfn11.

Furthermore, we observed a lack of correlation between the levels of Slfn11 and the anti-viral effect of this protein. Slfn11 affected HIV-1 and flavivirus infection in A172-SCR and −BC cells to a similar extent although these cells express markedly different levels of Slfn11. These observations indicate that above certain amounts of Slfn11 the anti-viral activity of this protein reaches saturation.

The proposed role of Slfn11 in maintaining the stability of the tRNA pool could also explain the direct correlation between the sensitivity of cells to genotoxic drugs and their Slfn11 levels. It has been found that cancer cells expressing higher levels of Slfn11 are more sensitive to DNA-damaging agents [30–34]. Then, it is possible that higher levels of Slfn11 could reduce the abundance of specific tRNAs in the cell affecting protein synthesis, as recently reported [3]. This effect will down-regulate the levels of DNA repair proteins encoded by codon-biased open reading frames [35], increasing the susceptibility of cancer cells to genotoxic agents.

In summary, our data indicate that Slfn11 opposes virus-induced changes in the tRNA repertoire, thus affecting evolutionarily unrelated viruses that manipulate the host tRNA repertoire to facilitate viral replication.

## MATERIALS AND METHODS

### Cell and Virus Culture

HEK293T, HeLa, SupT1, LLC-MK2, BHK-21, and A172 cells were obtained from the American Type Culture Collection (Manassas, VA). HEK 293T, HeLa, BHK-21, and A172 cells were maintained in Dulbecco’s Modified Eagle’s (DMEM) medium, LLC-MK2 and BHK-21 cells in Eagle’s Minimum Essential Medium (E-MEM), and SupT1 cells in RPMI 1640. These culture media were supplemented with 10% Fetal Bovine Serum (FBS) and 1% penicillin, streptomycin, and 1% non-essential amino acids (NEAA), 1% sodium pyruvate. Maintenance media used to perform viral infections consisted of E-MEM w/L-glutamine, supplemented with 2% FBS, 1% NEAA, 1% sodium pyruvate and 1% penicillin and streptomycin.

The WNV strain TVP-7767 (Passage: Vero, #3), RVFV strain MP-12 (Passage: Vero, #3) and ZIKV strain MR-766 (Passage: suckling mice brain, #150. Vero cells, #3) were obtained from the World Reference Center for Emerging Viruses and Arboviruses, University of Texas Medical Branch. DENV-2 strain 16681 (Passage: C6/36, #9) was obtained from the Navy Medical Research Center-6. VSV engineered to express eGFP has been previously described. Viral stocks were prepared in Vero cells maintained in E-MEM supplemented with 2% FBS. Titer of each virus stock was determined via plaque assay as described below.

A replication-defective HIV-1 reporter virus (Hluc) was used that expressed LTR-driven luciferase from the NEF slot and contains a large deletion in ENV[22]. Hluc was generated by calcium phosphate transfection of the corresponding HIV-1 expression plasmid (pHluc, 15 ug) and the VSV-G encoding plasmid pMD.G (5 ug) into HEK293T cells, as described before[22]. In accordance with World Health Organization’s and the Centers for Disease Control and Prevention’s guidelines, all work involving infectious WNV was performed in a biosafety level (BSL)-3 laboratory in accordance to biosafety practices described in the May, 2018 revised version of the University of Texas at El Paso’s (UTEP) BSL 3 Biological Safety Manual and Standard Operating Procedures. All work involving DENV, VSV, ZIKV, HIV-1, and RVFV MP-12 was performed in BSL-2+laboratory in accordance to biosafety practices described in the UTEP Biological Safety Manual.

### Virus replication dynamics

All cell lines infected with WNV, DENV, RVFV, ZIKV and VSV were seeded in T25 cell culture flasks (2.5×10^5^ cells in 2ml total volume) and allowed to grow overnight. The following day the cells were infected with respective viruses and incubated at 37°C for 1 h. Cells were subsequently washed three times with serum-free medium to remove input virus, replenished with maintenance medium and incubated at 37°C. Cell culture supernatants were collected every 8 hrs until experiments were stopped and stored at −80°C.

### Plaque Assay

Cell supernatants containing WNV or VSV were subjected to ten-fold serial dilutions, and inoculated onto confluent monolayers of LLC-MK2 cells in 12-well cell culture plates and incubated at 37°C for 1 h with gentle rocking every 15 minutes. The cells were then overlaid with 1 ml of 0.5% agarose in E-MEM maintenance medium. Cells were incubated at 37°C for 3 days and then stained with 1g/L of Napthol Blue Black, 13.6g/L of Sodium Acetate Anhydrous, 60ml/L glacial acetic acid to visualize plaques. Plaque formation on each cell line was quantified and viral titers were expressed as plaque-forming units per milliliter (PFU/ml).

DENV titers were determined as previously described[36]. Briefly, 3 × 10^5^ BHK-21 cells were seeded in 12-well cell culture plates, and then infected with viral supernatants at 37°C for 3 hrs, followed by the addition of 1ml of 3% carboxymethylcellulose overlay medium. Cells were cultured for 5 days, followed by staining and quantification as described above.

RVFV and ZIKV plaque assays were performed as described for WNV but using Vero 76 cells. RVFV- and ZIKV-infected cells were cultured at 37°C for 4 or 5 days, respectively. Staining and quantification was performed as described above.

Plaque assays for estimating viral titers were conducted in triplicate experiments with samples derived from independent viral infections.

### Immunoblotting

Full procedures for protein detection by immunoblot have been described previously[37]. Briefly, cellular lysates were obtained by lysing cells with 2x Laemmli Buffer and boiling for 10 minutes. Cell lysates were resolved by SDS-PAGE and transferred overnight to PDVF membranes at 100 mA at 4°C. Membranes were blocked in TBS containing 10% milk for one hour and then incubated in the corresponding primary antibody diluted in TBS-5% milk-0.05% Tween 20 (antibody dilution buffer). Full-length Slfn11 and Slfn11 C-terminus mutant were detected with anti-Slfn11 antibody E-4 (Santa Cruz Biotechnology 1/500). The Slfn11 N-terminus mutant was detected with anti-Slfn11 antibody D-2 (Santa Cruz Biotechnology 1/500). WNV envelope protein was detected with antibody PA1-41073 (Thermo Fisher Scientific, 1/500). Tetherin (BST-2) was detected with anti-BST-2 antibody (Santa Cruz Biotechnology 1/500). α-tubulin was detected as a loading control with antibody from clone B-5- 1-2 (Sigma, 1/4000). Membranes were incubated overnight at 4°C with primary antibodies, whereas anti-α-tubulin Mab was incubated for 30 minutes at 25°C. Primary antibody-bound membranes were washed in TBS-0.1% Tween 20 and bound antibodies detected with goat anti-mouse IgG-HRP (1/2000, Sigma) or a goat anti-rabbit IgG-HRP (1:4000, EMD Millipore) followed by chemiluminescence detection. Densitometry analysis of selected bands was quantified based on their relative intensities using Image Studio Software (LI-COR, Lincoln, NE).

### Plasmids

For the generation of HIV-1-derived viral vectors, plasmids were obtained from Eric Poeschla laboratory (Mayo Clinic, Rochester, MN) ^10^. These lentiviral vectors were used to express Slfn11- and control-shRNAs and Slfn11 proteins. They were generated with packaging plasmid pCMV∆R8.91, a transfer plasmid derived from pTRIP (described below), and the envelope plasmid pMD.G encoding the vesicular stomatitis virus glycoprotein G (VSV-G).

Slfn11- and SCR-shRNA plasmids: An shRNA construct (Top: 5’- GATCCGGCTCAGAATTTCCGTACTGAATTCAAGAGATTCAGTACGGAAATTCTGAGCTTTTTTGGAAA-3’, Bottom:5’-AGCTTTTCCAAAAAAGCTCAGAATTTCCGTACTGAATCTCTTGAATTCAGTACGGAAATTCTGAGCCG-3’) against Slfn11 was designed using a target sequence that has been previously described[4]. Briefly, Slfn11 shRNA construct was ligated into the pSilencer 2.1 U6 Hygro shuttle vector (AM5760, Thermo Fisher Scientific) and sequence verified. Control shRNA contains a scrambled sequence (SCR) that was obtained from the negative control plasmid provided with the kit. The Slfn11 and SCR shRNA expression cassettes were amplified by PCR and ligated into a unique PpUMI site in the HIV-1-derived transfer plasmid pTRIP-eGFP and their sequences verified.

shRNA-resistant Slfn11 expression plasmid: The shRNA-resistant Slfn11 cDNA was engineered by introducing 7 synonymous mutations within the 21nt-long shRNA target sequence of Slfn11. Plasmid pCDNA-V5-His-Slfn11 (Michael David, University of California San Diego)[4] was used as template for the Phusion High-Fidelity DNA polymerase (ThermoFisher, F530S). Primers used to introduce mutations were forward, 5’-TCGGACCGAGGATGGGGACTGGTATGGG-3’ and reverse, 5’-AAGTTTTGCGCTTCGTCAATGACG-3’. The newly created shRNA escape mutant cDNA was then amplified using the high-fidelity Deep Vent DNA polymerase (New England Biolabs), digested with SbfI and SpeI restriction enzymes, cloned into unique SbfI-SpeI sites in the pTRIP-IRES-P HIV-1-derived transfer plasmid, and the sequence verified.

N- and C-terminal Slfn11 mutant expression plasmids: pTRIP-IRES-P-Slfn11-shRNA-resistant plasmid was used as template to generate the Slfn11 truncated mutants using the QuikChange Lightning site-directed mutagenesis kit (Agilent Technologies). The mutant expressing N-terminal Slfn11 (amino acids 1-441) was generated with primers forward 5’-GAACAAAAACTCATCTCAGAAGAGGATCTG-3’, and reverse 5’-GAAGATCAAAATTCCCCGAAAGAAAG-3’ whereas primers forward 5’-TCTAGAAGTTGGGCTGTGGACC-3’ and reverse 5’-CATACTAGTGGATCCTCTAGC-3’ were used to produce C-terminal Slfn11 (amino acids 442-901). Mutants were verified by DNA sequencing.

### Production of lentiviral vectors

The full procedures for transfection and production of lentiviral vectors has been described previously[22, 37, 38]. Briefly, HEK293T cells were calcium-phosphate transfected with the corresponding transfer plasmid derived from pTRIP (15 ug), the packaging plasmid pCMV∆R8.91 (15 ug), and VSV-G envelope expression plasmid pMD.G (5 ug). The viral supernatants were harvested 48 hr post-transfection and concentrated by ultracentrifugation at 124,750g for two hr on a 20% sucrose cushion.

Expression of Slfn11- and SCR-shRNA in A172 cells. A172 cells were transduced with shRNA-, eGFP-expressing lentiviral vectors and cells expressing the highest 10% of eGFP fluorescence were isolated by cell sorting and expanded in culture. Slfn11 levels were determined in these cells by immunoblot, as described above.

Expression of Slfn11 full-length and deletion mutants in Slfn11-dificent cell lines: A172-KD, HEK293T, BHK-21, and HeLa cells were engineered to express Slfn11 proteins by transduction with lentiviral vectors expressing Slfn11 and the puromycin resistant gene. Briefly, viral vectors were produced in HEK293T by transfection with the transfer plasmid pTRIP-IRES-P-Slfn11-shRNA-resistant plasmid expressing Slfn11 full-length or deletion mutants (15 ug) and the packaging and envelope expression plasmids described above. Viral supernatant were concentrated by ultracentrifugation and used to transduce cells. Three days later, transduced cells were selected in the presence of puromycin (A172-KD and HEK293T: 3ug/ml, HeLa: .375ug/ml, BHK-21: 6ug/ml). Slfn11 expression was verified by immunoblot.

### Single-Round Infectivity Assay

HEK293T-, HeLa- and A172-derived cells were seeded onto 24-well plates (2×10^4^ cells/well) and allowed to grow overnight. Next day, cells were infected with Hluc and 24 hrs later the cells were extensively washed to remove the input virus. Four days later cell culture supernatant was collected for HIV-1 p24 quantification and cell lysates prepared in a buffer containing 1% Triton X-100 for luciferase activity quantification (Bright-Glow Luciferase Assay System, Promega), following the manufacturer’s instructions. Luciferase activity was determined in triplicate samples using a microplate luminometer reader (Thermo Scientific, Luminoskan Ascent). Luciferase and HIV-1 p24 samples were derived from at least three independent infections.

### HIV-1 p24, IFN-α and IFN-β ELISAs

HIV-1 infection was measured by quantifying HIV-1 p24 in the supernatant of infected cells (described above) by ELISA (ZeptoMetrix Corporation, 0801008). IFN-α and β was quantified in the cell supernatant of the infected cells by ELISA (PBL Assay Science, Cat # 41115-1 for IFN-α and # 41415-1 for IFN-β). ELISAs were performed according to the manufacturer’s instructions.

### Indirect immunofluorescence microscopy

This technique was used for determining WNV infection in A172-derived cells and for localizing Slfn11 in A172- and HEK293T-derived cells. A172-derived cells (1.5×10^4^/well) were seeded onto 96-well confocal microscopy plates and infected with WNV at MOI 1 twenty-four hours later. Infected cells were fixed and permeabilized with Cytofix/Cytoperm buffer (BD Biosciences, Cat#554714) at 24hrs and 48hrs post-infection and then stained with an anti-Flavivirus group antigen monoclonal antibody that recognizes WNV Env (ATCC, clone D1-4G2-4-15, 1/200) for 2hrs at room temperature. Cells were then washed 3 times with PBS containing 0.1% FBS and then incubated with Alexa 568-conjugated antibody (Invitrogen, H-11004, 10ug/ml) for 1h at room temperature. Cells were again washed 3 times with PBS and cell nuclei was stained with Hoechst 33342 (Invitrogen, H-3570, 20ug/ml) for 10 min.

For subcellular localization of Slfn11, uninfected A172- and HEK293T-derived cells were staining as described above. Specifically, Slfn11 full-length and C-terminus were detected with the anti-Slfn11 antibody E-4 (Santa Cruz Biotechnology, sc-374339, 1/200) and the Slfn11 N-terminus mutant was detected with anti-Slfn11 antibody D-2 (Santa Cruz Biotechnology sc-515071,1/200).

### tRNA PCR-array and analysis

To determine the effects of WNV infection on the host tRNA repertoire, we quantified the tRNA pool using a PCR-based methodology previously described[10]. The procedure and data analysis was performed by Arraystar Inc. using the nrStarTM Human tRNA Repertoire PCR Array. A172-derived cells were infected with WNV at MOI 1 for 1hr at 37°C washed with fresh culture media and replenished with culture media. 8 hrs post-infection, cells were collected and total cellular RNA was extracted using Trizol reagent. Experiments were performed in triplicate with appropriate non-infected controls. Total cellular RNA was sent to Arraystar Inc for analysis. Briefly, quality control was performed on extracted RNA samples by NanoDrop ND-1000 and RNA integrity and genomic DNA contamination were assessed by denaturing agarose gel electrophoresis. Next, RNA samples were subjected to a demethylation step, followed by first-strand cDNA synthesis (Arraystar, rtStarTM tRNA-optimized First-Strand cDNA Synthesis Kit, Cat# AS-FS-004). Real-time PCR was then performed using a proprietary human tRNA Repertoire PCR Array that is able to distinguish 66 nuclear tRNA isodecoders, covering all anti-codons available in GtRNAdb and tRNAdb databases. Three stably expressed small ncRNA genes RNU6-2 (Ref 1), SNORD43 (Ref 2), and SNORD95 (Ref 3) were included in the array as the quantification references for tRNA and data normalization. One external RNA Spike-In mix was added in the RNA sample prior to the first strand cDNA synthesis. The RNA Spike-In control assay indicated the overall success and the efficiency of the reaction beginning from the cDNA synthesis to the final qPCR. For positive PCR control, two replicates of one artificial DNA and the PCR primer pairs were used to indicate the qPCR amplification efficiency. Only Ct value greater than 35 were considered as good qPCR amplification efficiency and considered for analysis. The positive PCR control was used as an inter-plate calibrator and a control to exclude genomic DNA contamination.

### RNA extraction

Viral supernatants were treated with RNase (40ug/mL) and DNase (2U) for 30min at 37°C and then total RNA was isolated using Trizol LS reagent (Invitrogen, Cat#10296010) according to the manufacturer’s instructions. All RNA samples had absorbance ratios of 1.8–2.0 at 260/280 nm, indicating that samples were contaminant-free. Purified RNA samples were stored at −80°C until used.

### Quantification of WNV in cell culture supernatants

One-step RT-qPCR was carried out in the MiniOpticon™ Real-Time PCR System using the iTaq™ Universal SYBR^®^ Green One-Step RT-PCR Kit (cat #: 172-5151, Bio-Rad Laboratories, Hercules, CA, USA). After optimization, the reaction was performed in a final volume of 20 µl, including 10µl 2x SYBR® Green RT-PCR reaction mix, 0.6 µl (150 nM final concentration) of forward 5’-CCACCGGAAGTTGAGTAGACG-3’ and reverse 5’-TTTGCTAGCTTTAGGACCTACTATATCTACCTTTTGGTCACCCAGTCCTCCT-3’ primers, 0.25 µl iScript Reverse Transcriptase, and 5µl of RNA template. These primers are specific to the 3’UTR of the WNV genome and have been previously described[39]. The thermal cycling conditions consisted of a 10 min reverse transcription step at 50°C, 1 min of Taq at 95°C, followed by 45 cycles of PCR at 95°C of denaturing for 10 sec, 30 sec annealing at 65°C and 90 sec extension at 72.0°C with a step of a single fluorescence emission data collection followed by 10 min at 72°C for final extension. The specificity of amplicon was verified by melting curve analysis (55 to 95°C) with a heating rate 0.2°C per 5 sec to verify the identity and purity of the amplified product. Triplicate reactions were carried out for each sample, and no template control was included. Cycling conditions and annealing temperatures were optimized using RNA purified from a WNV stock.

### Bioinformatics analysis

Analysis of the impact of WNV-induced changes in tRNA on viral protein translation was performed by using a python script created to determine the differential tRNA expression values over the open reading frames within the viral genome. In short, the tRNA differential expression values were uploaded from text file into a python dictionary. For each codon not found within the tRNA expression dataset the codon value was set to 0, and for redundant codons it was limited to the value of the highest expression to limit the codon to a single value. Additionally, to make visualization of the values easier, a sliding window approach was taken to average the differential expression over a desired number of amino acids. The window was then shifted by a consistent step size, and the average again determined. To allow for variation in the resolution of the graphs, the window size and step size were coded as adjustable parameters within the python script for easy adjustment. For each gene, the individual codon usage and the windowed average values were written to a CSV file. The CSV file was then imported into Excel and graphs were then created from the windowed codon usage.

### Statistical analysis

All data used for viral replication curves were transformed to log_10_pfu/ml. Repeated-measures ANOVA [74] was used to test the impact of different cell lines expressing or not Slfn11 on viral replication curves and the Tukey-Kramer post-hoc test was used to identify significant differences in viral titer between cell lines.

## ACKNOWLEDGMENTS

We thank Dr. Zachary Martinez [University of Texas at El Paso (UTEP)] for help with HIV-1 p24 ELISAs, and Dr. Michael David (University of California San Diego) for providing us with a Slfn11 expression plasmid.

